# Increased interaction between connexin43 and microtubules is critical for glioblastoma stem-like cell maintenance and tumorigenicity

**DOI:** 10.1101/2024.01.26.576347

**Authors:** James W. Smyth, Sujuan Guo, Lorie O’Rourke, Stacie Deaver, Jacob Dahlka, Elmar Nurmemmedov, Zhi Sheng, Robert G. Gourdie, Samy Lamouille

## Abstract

Glioblastoma (GBM) is the most common primary tumor of the central nervous system. One major challenge in GBM treatment is the resistance to chemotherapy and radiotherapy observed in subpopulations of cancer cells, including GBM stem-like cells (GSCs). These cells hold the ability to self-renew or differentiate following treatment, participating in tumor recurrence. The gap junction protein connexin43 (Cx43) has complex roles in oncogenesis and we have previously demonstrated an association between Cx43 and GBM chemotherapy resistance. Here, we report, for the first time, increased direct interaction between non-junctional Cx43 with microtubules in the cytoplasm of GSCs. We hypothesize that non-junctional Cx43/microtubule complexing is critical for GSC maintenance and survival and sought to specifically disrupt this interaction while maintaining other Cx43 functions, such as gap junction formation. Using a Cx43 mimetic peptide of the carboxyl terminal tubulin-binding domain of Cx43 (JM2), we successfully ablated Cx43 interaction with microtubules in GSCs. Importantly, administration of JM2 significantly decreased GSC survival *in vitro*, and limited GSC-derived tumor growth *in vivo*. Together, these results identify JM2 as a novel peptide drug to ablate GSCs in GBM treatment.

## Introduction

Glioblastoma (GBM) is a highly malignant and lethal cancer of the central nervous system. The current multimodal therapy for newly diagnosed GBM patients includes surgical resection, radiotherapy, and chemotherapy with temozolomide (TMZ), conferring a median survival of only 14.6 months^1-3^. Failure to generate more effective treatment strategies is due to the infiltrative nature of GBM tumor cells preventing complete surgical resection^4^, and the cellular heterogeneity within GBM tumors, which comprise a sub-population of GBM stem-like cells (GSCs) that are resistant to chemotherapeutic agents including TMZ^5-8^. In fact, GSCs present a high degree of plasticity with the ability to self-renew through symmetric proliferation or differentiate through asymmetric division, recapitulating the heterogeneous tumor and overall promoting GBM recurrence^9-11^. Multiple signaling pathways, including Notch, participate in the generation and maintenance of GSCs^12,13^. Notch signaling occurs via activation by single-pass ligand proteins present on the membrane of adjacent cells, with Notch receptors being cleaved and an intracellular domain translocating to the nucleus to activate the transcription of genes including the primary targets of Notch signaling, *HES* and *HEY*. Hes and Hey are transcriptional repressors that, among other functions, contribute to the self-renewal of GSCs^12,14-16^.

Intercellular junctions are crucial to maintaining homeostasis and provide mechanical communication between cells as well as, in the case of gap junctions, direct coupling of neighboring cellular cytoplasms. Dysregulation of these junctions is associated with numerous disease processes, and is critical to carcinogenesis particularly through facilitating invasion and cancer cell spread^17^. The gap junction protein Connexin43 (Cx43) is understood to be both tumor suppressive and oncogenic, depending on the stage/phase of cancer^18^. Recent studies have shown that increased Cx43 levels correlate with TMZ resistance in GBM cells^19-21^. In addition, brain metastatic cells utilize Cx43 to communicate with normal astrocytes to support tumor growth, invasion, and chemoresistance^22^. Importantly, Cx43 has also been associated with anti-proliferative effects in glioma and reduced levels of Cx43 protein was reported in high-grade gliomas, which highlights a complex dual role for Cx43 in GBM^23,24^.

Cx43 is a four-transmembrane protein that oligomerizes to form connexon hemichannels at the trans-Golgi network before trafficking to the plasma membrane through vesicular transport along microtubules. Once at the cell surface, connexons on apposing cells couple to form gap junction channels that cluster together and allow the passage of small molecules (<1 kDa), including ions and several second messengers^25^. Cx43 is associated with regulation of a variety of cellular functions, which it effects through channel -dependent and -independent mechanisms; including regulation of cell proliferation, migration/invasion, and apoptosis^18^. Therefore, detailed analysis of the subcellular localization of Cx43, rather than its expression alone, is critical in understanding the relationship between Cx43 and GBM progression.

Low levels of Cx43 expression and formation of associated gap junctions have been observed in GSCs in previous studies^26,27^. Given the significant non-junctional roles of Cx43 associated with cancer progression, we sought to isolate the function of Cx43 in these GSCs that reportedly have few gap junctions. Using super-resolution Stochastic Optical Reconstruction Microscopy (STORM), we primarily observed intracellular Cx43 decorating microtubules in GSCs, demonstrating for the first time such clustering *in situ*. The Cx43 protein harbors multiple protein-protein interaction motifs within its cytosolic carboxy-terminus (CT), including a tubulin binding domain^23,28^. To test the role of Cx43 interacting with microtubules in GSCs, we utilized a Cx43 mimetic peptide named JM2 (juxta<u>m</u>embrane 2) composed of the Cx43 CT amino acids 231–245 encompassing the microtubule binding sequence, and an antennapedia cell penetration domain that promotes cellular uptake. Our data show that JM2 efficiently disrupts interaction between Cx43 and microtubules, and significantly decreases TMZ resistant GSC survival *in vitro* and GSC-derived tumor growth *in vivo*. Together, these results identify a novel tumorigenic channel-independent role of Cx43 in GSCs through direct interaction with microtubules. JM2 could represent a novel therapeutic peptide specifically modulating Cx43 tumorigenic function to eradicate TMZ-resistant GSCs and improve GBM treatment through delaying tumor recurrence.

## Material and Methods

### Cell culture

Human glioblastoma stem cell (GSC) lines LN-229/GSC and primary GSC lines VTC-001, VTC-034, and VTC-037 were previously isolated and established^21,29,30^. GSCs were maintained in serum-free stem cell media of DMEM with high glucose and L-glutamine (Genesee Scientific), containing Gibco B-27 supplement (Thermo Fisher), fibroblast growth factor-2 (20 ng/mL; PeproTech), epidermal growth factor (20 ng/mL; PeproTech), penicillin (100 ug/mL) / streptomycin (100 IU/mL; Genesee Scientific), and MycoZAP Plus-PR (Lonza). GSCs were passaged using TrypLE Express (Gibco). To induce GSC differentiation, medium was changed to DMEM with high glucose and L-glutamine, penicillin (100 ug/mL), streptomycin (100 IU/mL), MycoZAP Plus-PR, and 10% Fetal Bovine Serum (Gibco). Normal human astrocytes (NHA) were purchased from Lonza, and cultured in AGM™ Astrocytes Growth Medium BulletKit™ (Lonza). NHA were passaged using Trypsin 0.25% (Quality Biological) after one wash with Phosphate Buffer Saline (PBS; Genesee Scientific) without Ca^++^ and Mg^++^.

### Peptide and temozolomide treatment

Lyophilized peptides were obtained from Peptron, Inc. (South Korea) with purity > 95%, reconstituted in water or in PBS at a concentration of 10 mM, and aliquots were stored at −80°C freezer. Reconstituted peptides were added to the cell culture medium at increasing concentrations for times indicated. Temozolomide (TMZ; Selleck Chemicals) was reconstituted in DMSO at a concentration of 50 mM and aliquots were stored at −20°C freezer. TMZ was directly added to the cell culture medium at different concentrations for times indicated.

### Gliosphere formation assay

GSCs were plated as single cell suspension in low-attachment 96-well plate (100-500 cells per well), using methods similar to those reported previously^21,29^. Cells were then treated with peptides every other day. 2–3 weeks later, gliospheres were observed using phase contrast microscopy on a Revolve microscope (Echo Laboratories), and the number of gliospheres was determined.

### MTS viability assay

GSCs were plated in a 96-well plate (2,000 – 5,000 cells per well). Cells were then treated with TMZ (10, 50, or 100 μM), antennapedia, JM2-scrambled, or JM2 (10, 50, or 100 μM). After 4 days, cell viability was monitored following addition of MTS reagent (Promega) according to manufacturer’s instructions. The absorbance at 490 nm was measured using a SpectraMax i3 or iD3 microplate reader (Molecular Devices).

### Caspase 3/7 activity assay

GSCs were plated in a 96-well plate (2,000 cells per well) and treated with TMZ or peptides as described above. The activity of caspase 3/7 was measured after 48 h using the Caspase-Glo® 3/7 assay (Promega) following the manufacturer’s instructions. Fold change of caspase 3/7 activity was defined as the ratio of caspase3/7 luminescence in treated cells compared to controls.

### Western blotting

Cells were lysed in RIPA buffer (0.1% sodium dodecyl sulphate, 50 mM Tris pH 7.4, 150 mM NaCl, 1 mM EDTA, 1% Triton X-100, 1% deoxycholic acid, 200 μM Na_3_VO_4_, and 1mM NaF) supplemented with HALT^TM^ Protease and Phosphatase Inhibitor Cocktail (Thermo Fisher). Lysates were sonicated prior to centrifugation at 4°C for 20 min at 13,000 rpm. Protein concentration was quantified using the DC protein assay (Bio-Rad Laboratories) and a standard curve obtained with increasing concentrations of Bovine Serum Albumin (BSA; Fisher Scientific), and protein samples were normalized to 20 μg total protein per lane resolved by SDS-PAGE electrophoresis. 4X Bolt LDS sample buffer supplemented with 400 mM DTT (final concentration 1X Bolt LDS with 100 mM DTT) was added to samples before heating at 70°C for 10 min. SDS-PAGE was run using NuPAGE Bis-Tris 4%-12% gradient gels (Thermo Fisher) with MES or MOPS running buffer according to manufacturer’s instructions. Spectra™ Multicolor Protein Ladder (Thermo Fisher) or Precision Plus Protein™ Kaleidoscope™ Prestained Protein Standards (Bio-Rad Laboratories) were included as protein ladders on the same gels. Proteins were transferred to a PVDF membrane using the Bio-Rad Turbo Transblot System and transfer kit (Bio-Rad Laboratories) and fixed in methanol prior to blocking in 5% non-fat milk in TNT buffer (50 mM Tris pH 8.0, 150 mM NaCl, 0.1% Tween 20) for 2 h at room temperature. Primary antibody labeling was performed overnight at 4°C using the following primary antibodies diluted in TNT buffer containing 3% BSA; rabbit anti-Cx43 (MilliporeSigma), rabbit anti-CD133 (Abcam), rabbit anti-Olig2 (MilliporeSigma), rabbit anti-Notch1 (Cell Signaling Technology), mouse anti-α-tubulin (MilliporeSigma), rabbit anti-β-tubulin (Abcam), rabbit or mouse anti-GAPDH (Santa Cruz Biotechnology). Membranes were washed three times prior to secondary antibody labeling for 1 h at room temperature with goat secondary antibodies against mouse or rabbit IgG conjugated to HRP (MilliporeSigma). Biotin-tagged peptides were labeled with Pierce™ High Sensitivity NeutrAvidin™-HRP (Thermo Fisher) directly after blocking. SuperSignal™ West Pico PLUS Chemiluminescent Substrate or SuperSignal™ West Femto Maximum Sensitivity Substrate (Thermo Fisher) were used to detect HRP-conjugated secondary antibodies according to manufacturer’s instructions prior to imaging. Membranes were imaged on a ChemiDoc™ imaging system (Bio-Rad Laboratories). When necessary, membranes were re-blotted using Re-Blot Plus Strong Solution (MilliporeSigma) for 1 h before blocking and immunoblotting with different primary antibodies.

### Co-immunoprecipitation

Prior to lysis, cross-linking was completed by incubating cells in DTBP (Dimethyl 3,3’-dithiobispropionimidate; 540 μg/mL in PBS) for 30 min at 37 °C followed by a glycine quench (100 mM glycine in PBS) for 15 min at room temperature. Cross-linked samples were lysed in RIPA buffer supplemented with HALT^TM^ Protease and Phosphatase Inhibitor Cocktail as described above, but without sonication. Co-immunoprecipitation (co-IP) assays were also performed without cross-linking; non-cross-linked samples were lysed using a co-IP lysis buffer (0.5% Triton X-100, 50 mM Tris HCl, 150 mM NaCl, 1 mM EDTA, 1 mM EGTA, 1 mM DTT, 0.1 mM Na_3_VO_4_, and 1 mM NaF) supplemented with HALT^TM^ Protease and Phosphatase Inhibitor Cocktail, without sonication. Protein concentration was quantified using the DC protein assay and a standard curve obtained with increasing concentrations of BSA, and 500 μg of total protein was used per reaction for co-IP. Inputs were removed prior to co-IP and denatured in Bolt LDS sample buffer as described above. Protein lysate was incubated with Protein G Sepharose^TM^ 4 Fast Flow beads (GE Healthcare) for pre-clearance for 30 min at 4°C. Samples were incubated with 2 μg of mouse anti-α-tubulin (MilliporeSigma) for cross-linked samples or mouse anti-Cx43 antibody (MilliporeSigma) for non-cross-linked samples for 90 min at 4 °C. Mouse-IgG (Fisher Scientific), antibodies against HA tag (Biolegend), or V5 tag (Cell Signaling Technology) were used as isotype controls. Samples were incubated with Protein G Sepharose^TM^ 4 Fast Flow beads for 1 h at 4 °C. Protein complexes were washed four times with RIPA or co-IP lysis buffers, then eluted and denatured in 2X Bolt LDS Buffer supplemented with 100 mM DTT. SDS-PAGE and western blotting occurred as described above using rabbit anti-Cx43 or rabbit anti-β-tubulin (MilliporeSigma) primary antibodies.

### Immunofluorescence

Cells were washed once with warm PBS before fixing in 4 % paraformaldehyde for 20 min at room temperature or ice-cold methanol for 5 min on ice, washed twice, and held in PBS at 4°C until immunostaining was conducted. Cells were permeabilized and blocked in 5% normal goat or donkey serum (Fisher Scientific) and 0.1% Triton X-100 in PBS for 2 h at room temperature. Primary antibodies rabbit anti-Cx43, mouse anti-pan Cadherin, or mouse anti-α-tubulin (MilliporeSigma) were diluted in blocking buffer and labeling was performed overnight at 4°C. Cells were washed six times with PBS (2 x quick, 2 x 10 min, 2 x 5 min) prior to incubating with goat anti-mouse or -rabbit IgG secondary antibodies conjugated to Alexafluor488, AlexaFluor555, or AlexaFluor647 (Thermo Fisher) for confocal; or donkey anti-mouse or -rabbit IgG secondary antibodies conjugated to CF568 (Biotium) or AlexaFluor647 (Jackson Immunoresearch) for STORM; for 1 h at room temperature. DAPI (Thermo Fisher) was included with secondary antibodies to counterstain nuclei for confocal only and was omitted from labeling for STORM. Membranes were labeled using AlexaFluor488- or AlexaFluor647-conjugated wheat germ agglutinin (WGA; Thermo Fisher). Slides were washed 6 times as before and mounted using Prolong Gold Antifade (Thermo Fisher) for confocal microscopy or maintained in PBS at 4°C prior to STORM imaging. To detect biotin-tagged peptides, fixed cells were incubated after the first set of washes with streptavidin conjugated to AlexaFluor647 (Thermo Fisher) diluted in high-salt buffer (0.5 M NaCl, 10 mM Hepes) for 30 min at room temperature. Cells were then washed four times with high-salt buffer (2 x quick, 2 x 10 min), and twice with PBS (2 x 5 min) before proceeding with secondary antibodies and washes as described above.

### Super-resolution (STORM) localization and analysis

Stochastic optical reconstruction microscopy (STORM) was conducted with a Vutara 350 microscope (Bruker). Immunolabeled cells were imaged in 50mM Tris-HCl, 10mM NaCl, 10% (wt/vol) glucose buffer containing 20 mM mercaptoethylamine, 1% (vol/vol) 2-mercaptoethanol, 168 active units/mL of glucose oxidase, and 1404 active units/mL catalase. 2,500 frames were acquired for each probe and 3D images were reconstructed in Vutara SRX software. Coordinates of localized molecules were used to calculate pair correlation functions in the Vutara SRX software.

### Cellular thermal shift assay (CETSA)

For an initial determination of the melting profile of α- and β- tubulin, fresh lysates of GSCs prepared in non-denaturing buffer was dispensed into 96-well PCR plate in stem cell medium (approximately 10,000 cells/well/50 µl), then was subjected to temperature gradient (37-65°C) for 20 min. Subsequently, centrifugation was performed at 14,000 rpm to sediment the unstable protein content. Supernatant was collected and SDS-PAGE gel was run, and immune-detection was performed to detect α- and β- tubulin using specific primary antibodies described above. Band intensities were quantified on LI-COR C-Digit Blot Scanner, and subsequently T_agg_(50) and T_agg_(75) values were calculated for α- and β- tubulin. In a subsequent run, fresh lysates of GSCs were treated at various doses with 3-fold dilutions (222.2, 74.0, 24.7, 8.2 and 2.7 nM) of peptides JM2 and JM2-scrambled, together with DMSO as control, for 1 h. Samples were then subjected to heat challenge at 57°C for 20 min, and unstable protein was removed by centrifugation step. Following an immunoblotting step, band intensities of the stable tubulin was quantified, and normalized to DMSO control. EC50 values of engagement for both peptide with α- and β- tubulin were subsequently calculated.

### Real-time quantitative PCR

RNA was extracted using the PureLink^TM^ RNA mini extraction kit (Thermo Fisher) and on-column DNA digestion was completed using PureLink^TM^ DNase according to the manufacturer’s instructions. RNA was reverse transcribed to generate cDNA using the iScript Reverse Transcriptions SuperMix for RT-qPCR kit (Bio-Rad Laboratories). Real-time PCR was performed using the SYBR select Master Mix for CFX (Thermo Fisher) in hard-shell 96-well PCR plates (Bio-Rad Laboratories) on a CFX Connect Real Time System (Bio-Rad Laboratories). Primer sequences used were: hNOTCH1_Fwd: 5’-CGCACAAGGTGTCTTCCAG-3’; hNOTCH1_Rev: 5’-CGGCGTGTGAGTTGATGA-3’; hHES1_Fwd: 5’-GGCTGGAGAGGCGGCTAA-3’; hHES1_Rev: 5’-GAGAGGTGGGTTGGGGAGTT-3’; hHEY1_Fwd: 5’-ACGAGACCGGATCAATAACA-3’; hHEY1_Rev: 5’-ATCCCAAACTCCGATAGTCC-3’.

### Animals

Animal studies were approved by the Institutional Animal Care and Use Committee (IACUC) of Virginia Tech. LN229/GCSs (1 × 10^5^) were mixed with Corning Matrigel® matrix growth factor reduced and subcutaneously injected into flanks of 8-week old male BALB/c Nude mice (Charles River Laboratories) as previously described^21,30^. After 7 days, tumors were measured and mice were separated into three groups; vehicle/control (PBS), JM2-scrambled, JM2. At day 9, vehicle or peptides were administered at 300 μM intratumorally every second day for 24 days. Tumor sizes were measured by electronic calipers at different times, and tumor volume was calculated using the formula (length×width^2^)/2.

### Tissue collection, sectioning, and staining

Tumors were harvested, snap-frozen in Tissue-Tek® O.C.T. compound (Sakura), and stored at - 80°C until sectioning. 10 μM sections were cut using a Leica CM1850 UV cryostat, placed onto Superfrost Plus^TM^ slides (Fisher Scientific) and fixed in -20°C acetone for 5 minutes before air drying. Slides were stored at -80°C prior to labeling. Tumors sections were then rehydrated in PBS for 5 min, permeabilized in 0.1% Triton X-100 in PBS for 1 h at room temperature, and washed four times with PBS (2 x quick, 2 x 5 min) prior to incubating with streptavidin conjugated to AlexaFluor647 (Thermo Fisher) diluted in high-salt buffer (0.5 M NaCl, 10 mM Hepes) for 30 min at room temperature. Tumor sections were then washed four times with high-salt buffer (2 x quick, 2 x 10 min), and twice with PBS (2 x 5 min) before using One-step TUNEL In Situ Apoptosis kit (Green Elab Fluor® 488; Elabscience), according to manufacturer’s instructions. Tumor sections were then washed in PBS (3 x 5 min), incubated with DAPI (Thermo Fisher) in PBS for 5 min, and washed again with PBS (4 x 5 min) before the slides were mounted using Prolong Gold Antifade (Thermo Fisher).

### Statistical Analyses

All quantification was performed on experiments repeated at least three times. Data are presented as mean ± SEM. Statistical analysis was conducted with GraphPad Prism 8 and 9 (GraphPad Software). Data were analyzed for significance using Student’s t test, one-way ANOVA with Tukey’s multiple comparisons test. A value of *p* < 0.05 was considered statistically significant. A within-between interaction repeated measure ANOVA was performed to evaluate differences in the rate of tumor growth among interventions. Specifically, value was the dependent variable and group, time, and a group by time interaction were the independent variables. A random effect was included for each mouse to handle the repeated measures.

## Results

### Cx43 displays increased cytosolic interaction with microtubules in glioblastoma stem-like cells

Primary human glioblastoma stem-like cells (GSCs) VTC-001, VTC-034, and VTC-037 previously isolated from freshly resected GBM tumors^21,29^ were cultured in stem cell medium as GSC-derived gliospheres or adherent GSCs, or differentiated by addition of 10% FBS (Figure 1A). Stemness was confirmed in gliospheres and adherent GSCs by western blotting, as observed by expression of GSC markers CD133 and Olig2 (Figure 1B). FBS-induced differentiation caused a decrease in CD133 and Olig2 expression after 24 h, which was further accentuated after 7 days (Figure 1B). Interestingly, we observed an increase in Cx43 expression at the protein level in differentiated VTC-001, VTC-034, and VTC-037 compared to their GSC counterparts (Figure 1C). Although Cx43 expression was lower in GSCs, confocal immunofluorescence microscopy revealed Cx43 primarily enriched within the cytoplasm of GSCs (Figure 1D).

**Figure 1:**
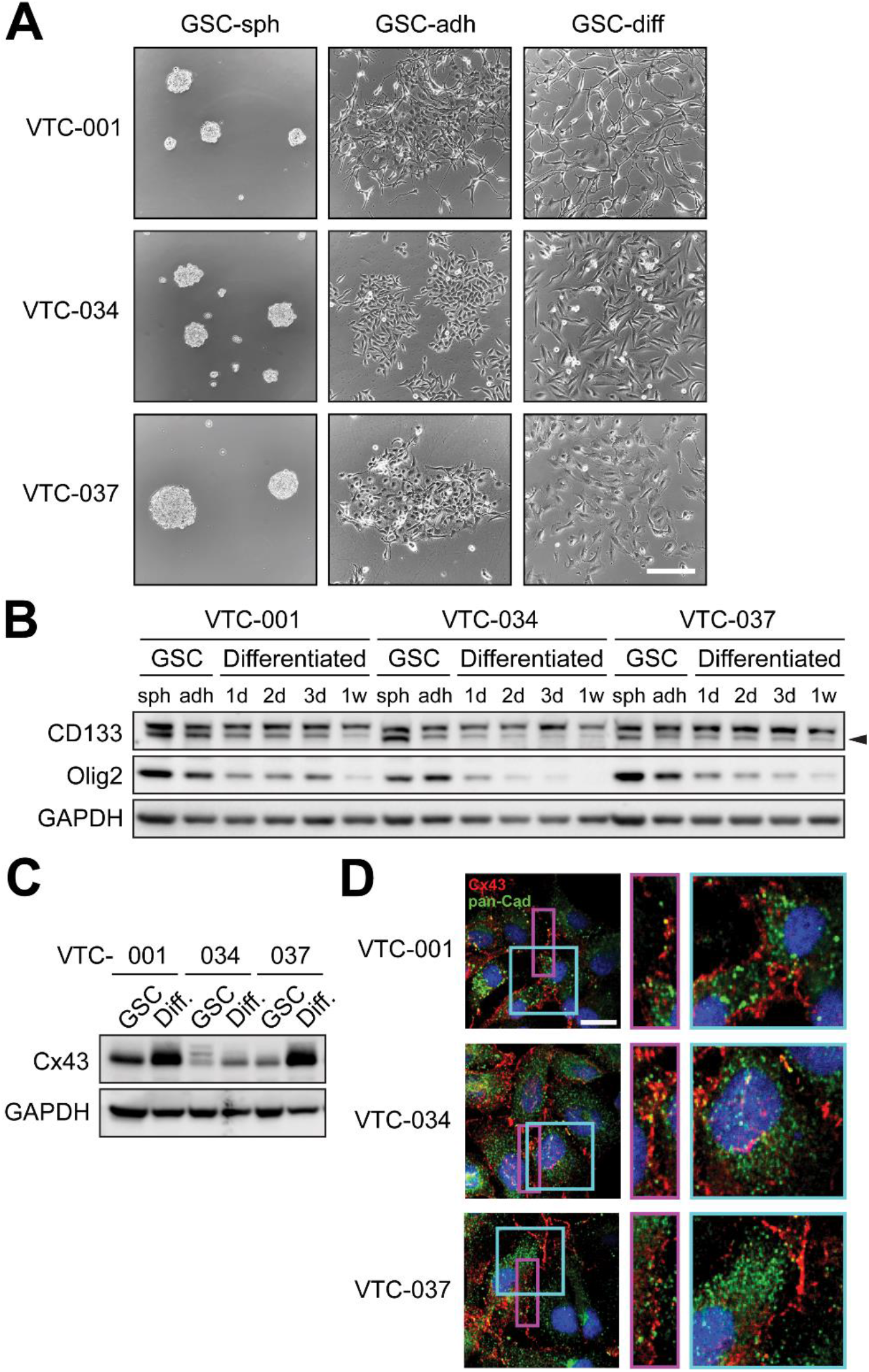
Cx43 expression and localization in GSCs. **A)** VTC-001, VTC-034, and VTC-037 GSCs were cultured in stem cell media as gliospheres or as adherent cells, or differentiated for 1 to 3 days in medium containing 10% FBS, and observed by phase-contrast microscopy. Scale bar: 200 μm. **B)** VTC-001, VTC-034, and VTC-037 cultured as GSC-derived gliospheres (sph), adherent GSCs (adh), or differentiated cells after addition of 10% FBS for 1, 2, 3, and 7 days, were lysed and analyzed by immunoblotting using antibodies against CD133, Olig2, and GAPDH as loading control. **C)** VTC-001, VTC-034, and VTC-037 GSCs were differentiated or not for 24 h, and cell lysates were analyzed by immunoblotting for Cx43, and GAPDH as loading control. **D)** VTC-001, VTC-034, and VTC-037 GSCs in culture were fixed and immunostained using antibodies against Cx43 (red), and pan-Cadherin (pan-Cad – green) to stain cell borders. Confocal immunofluorescence was used to observe Cx43 subcellular localization in the cytoplasm and at the cell borders. DAPI was used to stain nuclei. Scale bar: 20 μm.

We utilized STORM to further investigate the subcellular localization of Cx43 (green) in GSCs and found clear colocalization with the microtubule cytoskeleton (magenta) in VTC-001, VTC-034, and VTC-037 GSCs compared to differentiated GSCs (Figure 2A, Supplemental Figure 1A and B). Differentiating GSCs in medium containing 10% FBS resulted in a significant redistribution of Cx43 away from this direct interaction with microtubules (Figure 2A, Supplemental Figure 1A and B), quantified by pair correlation analysis (Figure 2B). These results were confirmed biochemically by co-IP, where again, decreased interaction between tubulin and Cx43 was observed following addition of 10% FBS in VTC-001 and VTC-037 GSCs (Figure 2C; lower band indicated by black arrow).

**Figure 2:**
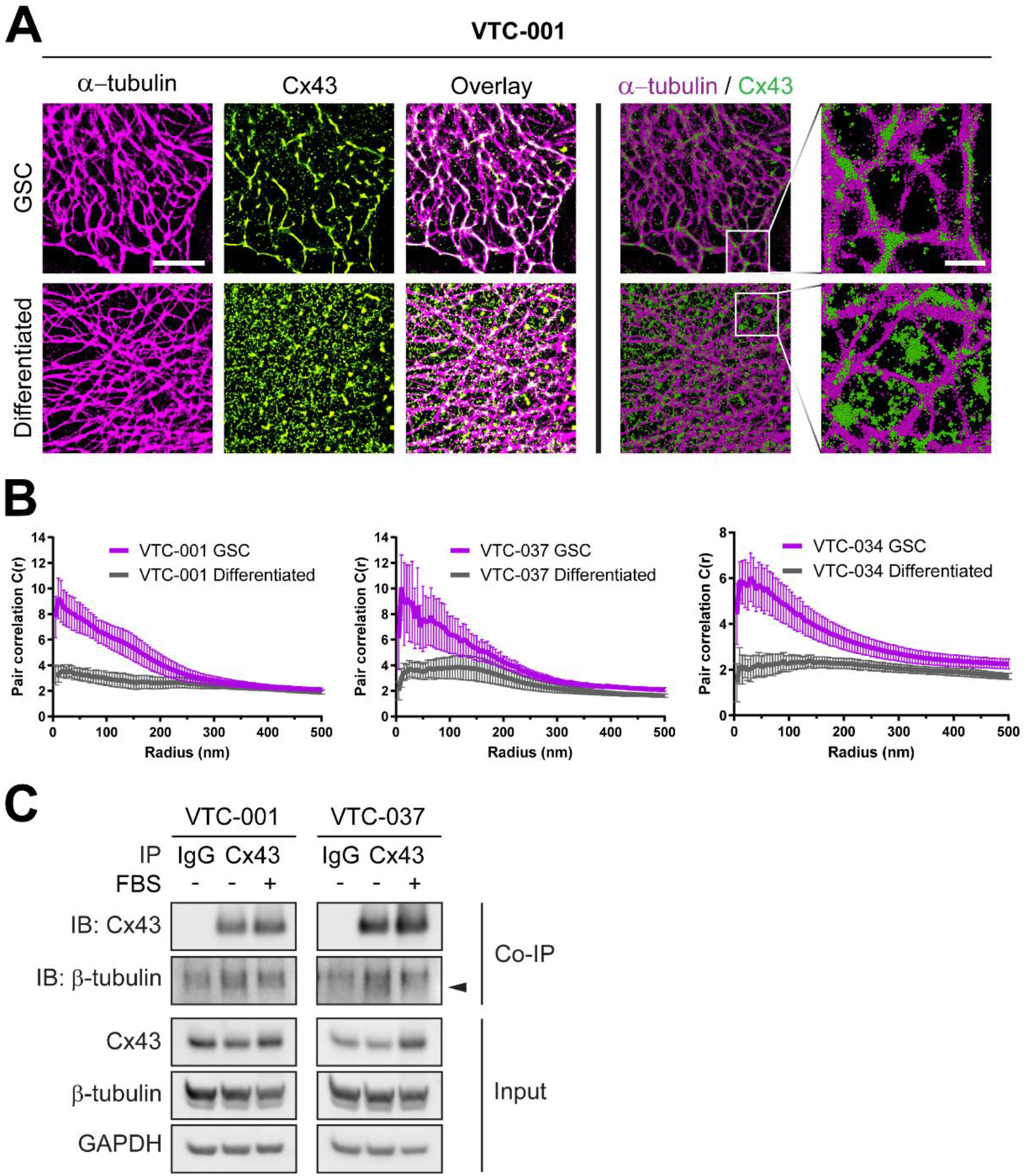
Increased Cx43 interaction with microtubules in GSCs. GSCs were differentiated through addition of FBS and interaction of Cx43 with the microtubule cytoskeleton was assessed by super-resolution microscopy and co-IP. **A)** Stochastic optical reconstruction microscopy (STORM) derived localizations of Cx43 (green) and α-tubulin (magenta) in VTC-001 GSCs or differentiated by addition of 10% FBS for 24 h. Left 6 panels - point- splatting is done to better identify co-localization (white; scale bar: 5 μm). Right 4 panels are point-clouds of 50 nm spheres representing individual localizations, including zoomed-in regions (scale bar: 1 μm). **B)** Cross-pair correlation functions for Cx43/α-tubulin complexing in VTC-001, VTC-034, and VTC-037 GSC and differentiated cell populations. (*n*=10). **C)** VTC-001 and VTC-037 GSCs were differentiated or not in 10% FBS and cell lysates were subjected to co-immunoprecipitation using Cx43 antibody or IgG for negative control, and/or immunoblotted using antibodies against Cx43, β-tubulin, and GAPDH for loading control. Black arrow indicates β-tubulin.

### Cx43 mimetic peptide JM2 disrupts Cx43 microtubule interaction with microtubules in GSCs

To assess the role of Cx43/microtubule interaction in GSCs, we utilized a Cx43 mimetic peptide, JM2, encompassing the Cx43 tubulin-binding domain, an antennapedia cell penetrating sequence, and a biotin tag for tracking, with biotin-tagged antennapedia and JM2-scrambled as negative controls (Figure 3A)^31^. Cellular uptake of JM2 at increasing concentrations was confirmed by western blotting in VTC-034 GSCs using HRP-conjugated streptavidin for detection (Figure 3B). JM2 cell uptake was apparent after 1 h, and sustained for 24 h before JM2 signal in GSCs decreased at 48 h (Figure 3C). Peptide uptake in GSCs was assessed by confocal immunofluorescence microscopy, where JM2 and JM2-scrambled (red) were confirmed to be localized within the cytoplasm of GSCs (Figure 3D). To further confirm that administered peptides were truly cytosolic, we performed Z-stack analyses using WGA (green) to identify post-Golgi cell membranes. 3D-projection of Z-stacks revealed enrichment of JM2 (red) within cells, as identified by localizations subjacent to WGA-labeled cell surfaces and colocalization with intracellular WGA signal (Figure 3E).

**Figure 3:**
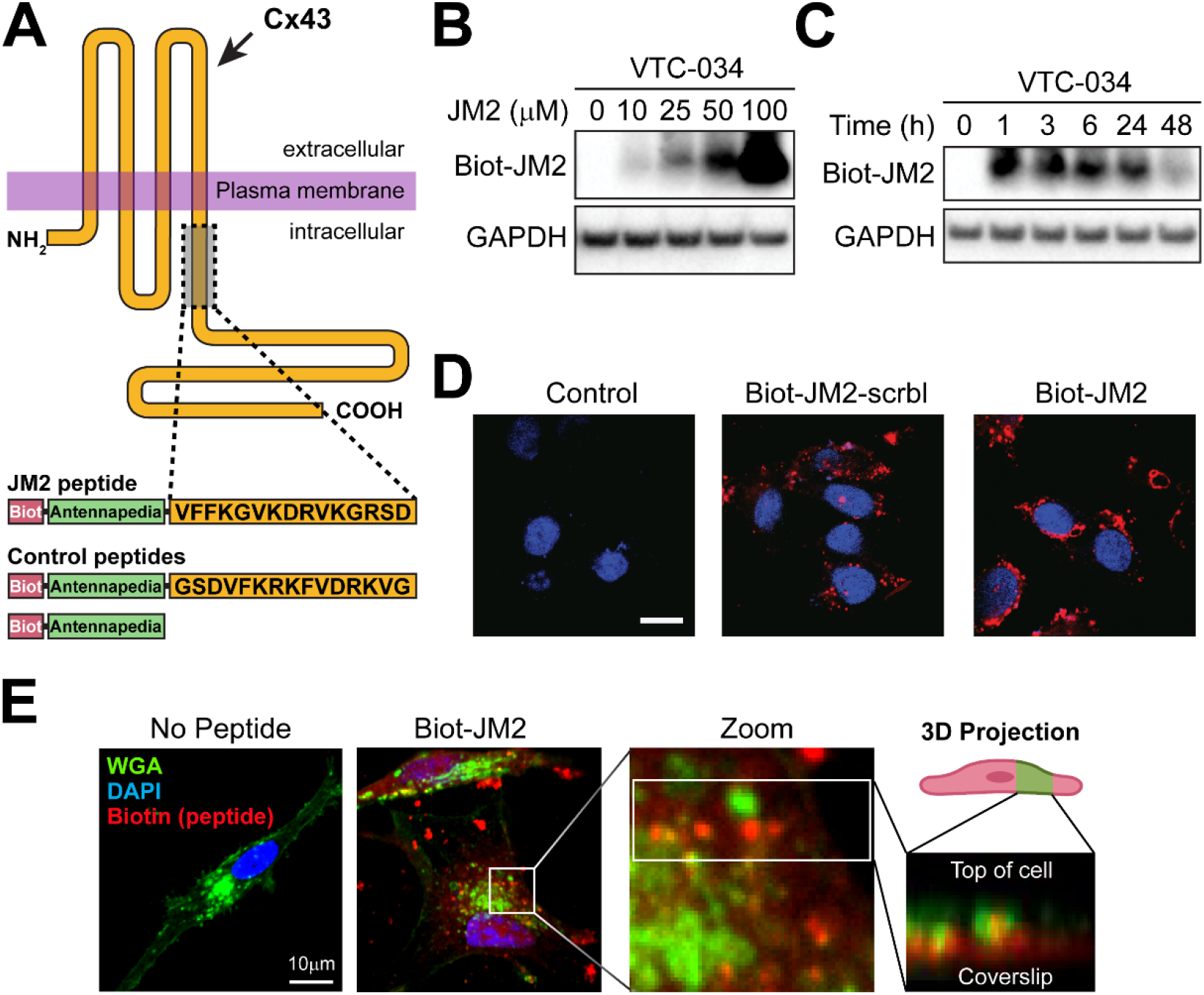
JM2 cell uptake in GSCs. **A)** Schematic of the JM2 peptide that encompasses the Cx43 tubulin-binding domain, an antennapedia cell penetration domain, and a biotin tag for tracking. Control peptides include JM2-scrambled and antennapedia. **B)** VTC-034 GSCs were treated or not with JM2 at different concentrations for 24 h, lysed and analyzed by electrophoresis using Neutravidin conjugated to HRP, and immunoblotting for GAPDH as loading control. **C)** VTC-034 GSCs were treated with JM2 at 50 μM for different times, lysed and analyzed by electrophoresis using Neutravidin conjugated to HRP, and immunoblotting for GAPDH as loading control. **D)** VTC-034 GSCs were treated or not with JM2-scrambled (JM2-scrbl) or JM2 at 50 μM for 24 h, and fixed. Biotin-tagged JM2-scrambled and JM2 were observed by confocal fluorescence microscopy using streptavidin conjugated to fluorophore (red), and DAPI was used to stain nuclei. Scale bar: 10 μm. **E)** Immunofluorescence confocal microscopy of VTC-037 GSCs treated or not with JM2 at 50 μM for 24 h before fixing, and probed for Wheat Germ Agglutinin (WGA – green), biotin-tagged JM2 (red), with nuclei identified using DAPI (blue). Original magnification: x100. Scale bar: 10 μm.

We next performed cellular thermal shift assay (CETSA) to characterize the specificity and affinity of JM2 interaction with α- and β- tubulin, i.e. primary microtubular subunits. In an initial heat gradient, thermal melting profiles of α- and β- tubulin were determined in VTC-037 GSC lysates with T_agg_(50) values of 49.5°C for α-tubulin, and 49.2°C for β-tubulin (Supplemental Figure 2). T_agg_(75) values for α- and β-tubulin were calculated as 57°C. Subsequent dose-gradients of the peptides JM2 and JM2-scrambled under T_agg_(75) revealed their selective target engagement potency (EC50) for α-tubulin and β-tubulin, while no EC50 was detected for actin, which was used as a negative control (Figure 4A). EC50 values for JM2 were calculated as 8.4 nM for α- tubulin, and 7.9 nM for β-tubulin, while EC50 values for JM2-scrambled were above 200 nM for both α-tubulin and β-tubulin (Figure 4B). These data confirmed that JM2 presented higher potency of target engagement with α-tubulin and β-tubulin compared to the JM2-scrambled control peptide. We then tested the ability for JM2 to disrupt Cx43 interaction with microtubules in GSCs utilizing STORM analysis, where we detected that JM2 significantly disrupts Cx43 interaction with microtubules in VTC-037 GSCs *in situ* (Figure C and D). These results were biochemically confirmed by co-immunoprecipitation experiments, wherein it was determined that JM2 robustly interacted with α-tubulin compared to antennapedia alone or JM2-scrambled peptides, concomitantly with a decrease in Cx43 interaction with α-tubulin (Figure 4E).

**Figure 4:**
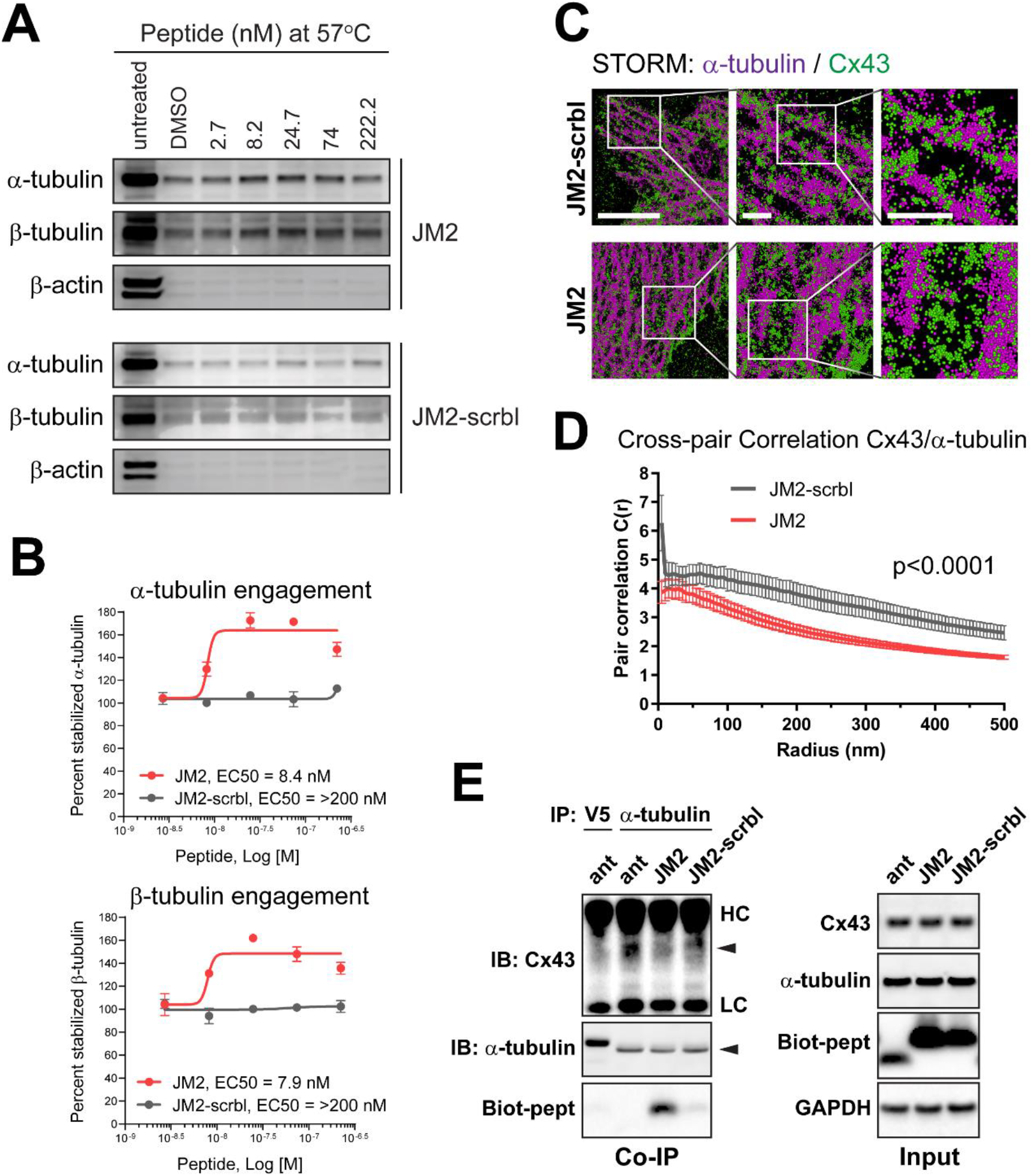
Cx43 mimetic peptide JM2 disrupts Cx43 – microtubule interaction. **A)** Cellular thermal shift assay in VTC-037 GSC lysates was used to determine JM2 and JM2-scrambled (JM2-scrbl) selective target engagement potency for α-tubulin and β-tubulin. VTC-037 GSC lysates were subjected to different concentration of JM2 or JM2-scrambled (JM2-scrbl) at 57°C, and peptide affinity was analyzed by western blotting using antibodies against α-tubulin, β-tubulin, and β-actin as negative control. **B)** Percentage of stabilized α-tubulin and β-tubulin at 57°C was represented, and EC50 values for JM2 and JM2-scrambled (JM2-scrbl) were calculated. **C)** STORM derived point-cloud localizations of Cx43 (green) and α-tubulin (magenta) in VTC-037 GSCs following treatment with JM2-scrambled (JM2-scrbl) or JM2 at 50 μM for 24 h. Zoomed out panels (left) scale bar: 6 μm. Zoomed in panels (middle) scale bar: 1 μm. Zoomed in panels (right) scale bar: 1 μm. Sphere size: 50 nm. **D)** Cross-Pair correlation functions for Cx43/α-tubulin interaction in C. (*n*=10). **E)** VTC-037 GSCs were treated or not with antennapedia (ant), JM2, or JM2-scrambled (JM2-scrbl) at 50 μM for 24 h. Following cross-linking, cell lysates were subjected to co-immunoprecipitation using α-tubulin antibody, or V5 antibody for negative control, and/or immunoblotted using antibodies against Cx43, α-tubulin, and GAPDH for loading control. Neutravidin-HRP was used to detect biotin-tagged peptides. HC: heavy chain; LC: light chain.

### JM2 peptide decreases GSC survival in vitro

We next determined the effect of JM2 in GSCs and observed that JM2 significantly decreased VTC-001, VTC-034, and VTC-037 GSC survival when cultured as adherent cells in a dose-dependent manner using an MTS assay (Figure 5A). As Cx43 is primarily expressed in astrocytes in the central nervous system^32^, we tested the effect of JM2 in these non-tumor cells. We did not observe a decrease in human astrocyte viability, even at higher JM2 concentrations (Figure 5B). As we previously reported a relationship between Cx43 and TMZ resistance, we tested the combination of TMZ and JM2 on GSC viability. While VTC-001 and VTC-037 GSCs are resistant to TMZ treatment, as previously described^21^, JM2 significantly decreased GSC viability as observed above, but the combination of TMZ with JM2 did not decrease GSC survival further (Figure 5C). These results were complemented by measuring caspase 3/7 activity as an indicator of apoptosis induction. We observed increasing doses of TMZ or JM2-scrambled were inefficient in inducing apoptosis in VTC-034 GSCs, whilst JM2 increased caspase 3/7 activity in these cells in a dose-dependent manner (Figure 5D). No effect on caspase 3/7 was observed in normal human astrocytes (Figure 5E). We next evaluated the effect of JM2 on GSC-dependent gliosphere formation, as an additional measure of GSC maintenance and survival. We found that JM2 inhibits VTC-001, VTC-034, and VTC-037 GSC-dependent gliosphere formation in a dose-dependent manner, with a concomitant significant decrease in gliosphere size and number following treatment (Figure 6A and B).

**Figure 5:**
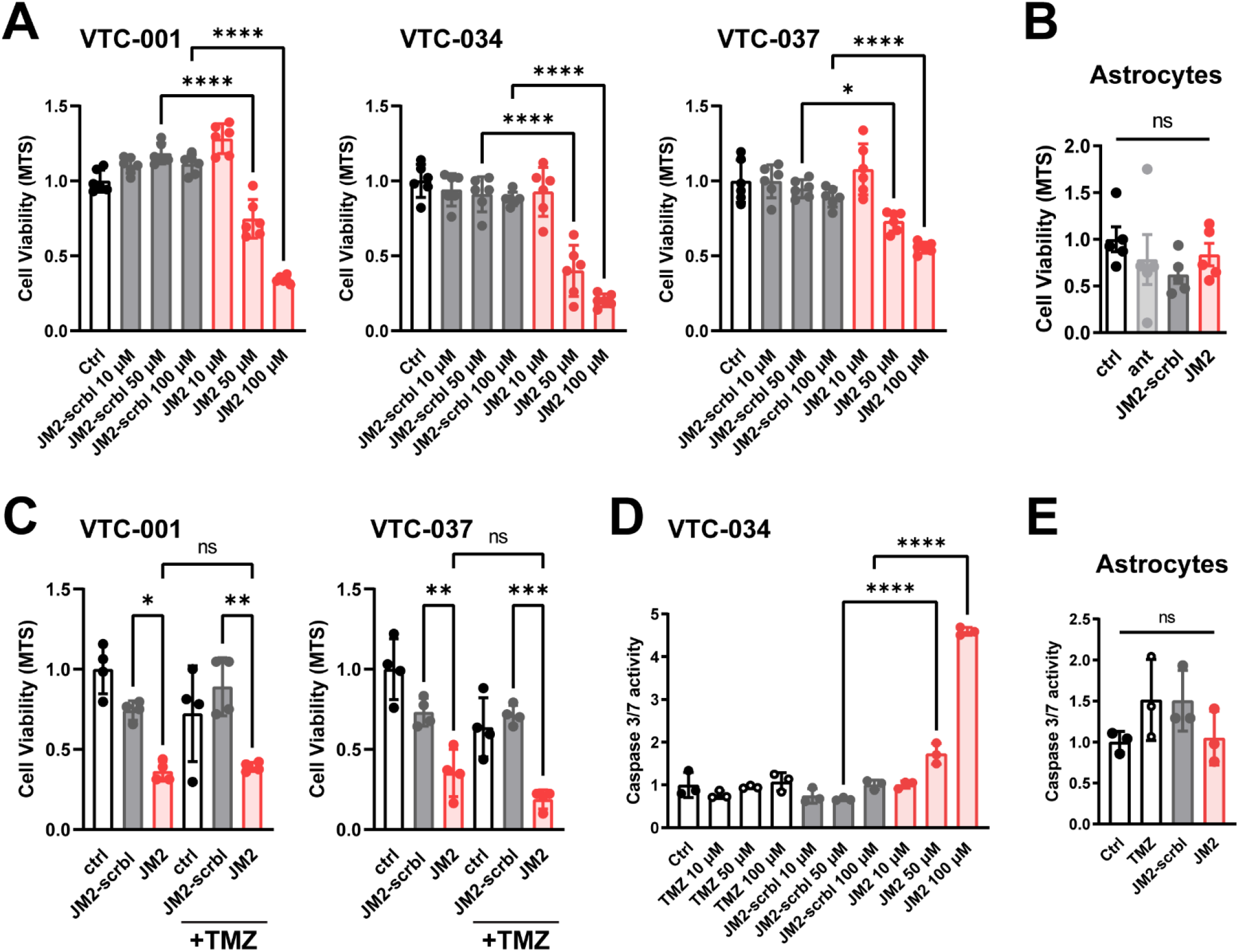
JM2 inhibits GSC survival *in vitro*. **A)** VTC-001, VTC-034, and VTC-037 GSCs were cultured as adherent cells in 96-well plates and treated or not with 10, 50 or 100 μM of JM2-scrambled (JM2-scrbl) or JM2 peptides for 4 days before cell survival was assessed using MTS assay. **B)** Human astrocytes were treated or not with 100 μM of antennapedia (ant), JM2-scrambled (JM2-scrbl) or JM2 peptides for 4 days before cell survival was assessed using MTS assay. **C)** VTC-001 and VTC-037 GSCs were cultured as adherent cells in 96-well plates and treated or not with temozolomide (TMZ) at 50 μM, with or without JM2-scrambled (JM2-scrbl) or JM2 peptides at 100 μM for 4 days before cell survival was assessed using MTS assay. **D)** VTC-034 GSCs were cultured as adherent cells and treated or not with different concentrations of TMZ, JM2-scrambled (JM2-scrbl), or JM2 for 4 days at 10, 50 or 100 μM before assessing apoptosis using caspase 3/7 assay. **E)** Human astrocytes were treated or not with 100 μM of TMZ, JM2-scrambled (JM2-scrbl) or JM2 for 4 days before assessing apoptosis using caspase 3/7 assay. Statistical analysis was performed with one-way analysis of variance (ANOVA) with Tukey’s multiple comparisons test. \**p*≤0.05, \*\**p*≤0.01, \*\*\**p*≤0.001, \*\*\*\**p*<0.0001. Data are represented as mean ± SEM.

**Figure 6:**
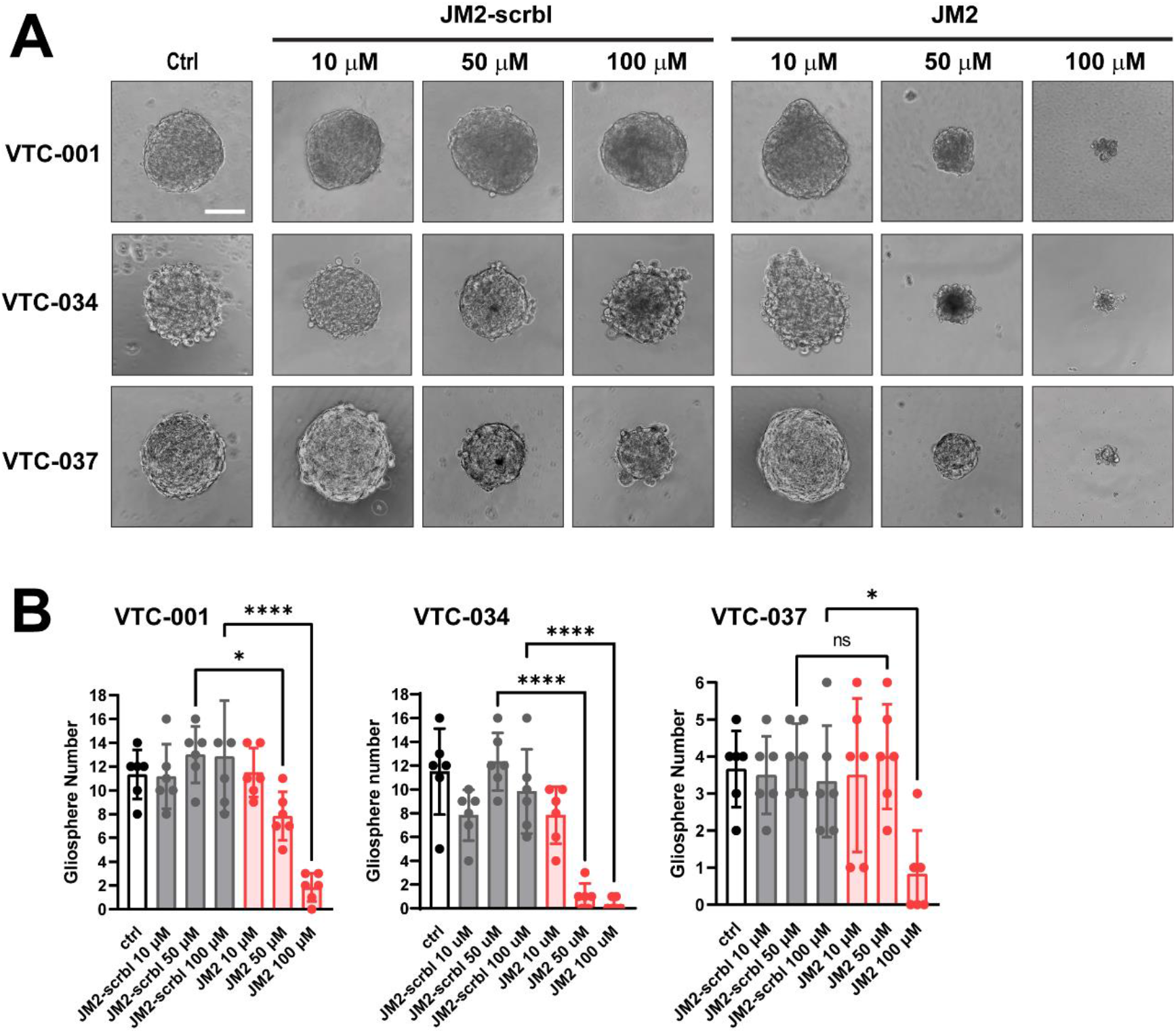
JM2 inhibits GSC-dependent gliosphere formation. **A)** VTC-001, VTC-034, and VTC-037 GSCs were cultured as single cells in suspension in low attachment 96-well plates, and treated or not with 10, 50 or 100 μM of JM2-scrambled (JM2-scrbl) or JM2 peptides every other day for 2 weeks. Gliospheres were observed by phase-contrast microscopy. Scale bar: 100 μm. **B)** Quantification of gliosphere numbers in A. Statistical analysis was performed with one-way analysis of variance (ANOVA) with Tukey’s multiple comparisons test. \**p*≤0.05, \*\*\*\**p*<0.0001. Data are represented as mean ± SEM.

### JM2 peptide inhibits Notch signaling

Notch signaling plays a critical role in GSC survival^12,14,33^. We tested the effect of JM2 on Notch1 expression and downstream targets in GSCs. JM2 significantly decreased Notch1 expression at the protein level after 24 h of treatment in VTC-001 and VTC-037 GSCs (Figure 7A and B), without affecting Cx43 expression (Figure 7C and D). Interestingly, Notch1 expression was not affected at the transcription level in VTC-037 GSCs (Figure 7E). Activation of Notch signaling results in Notch cleavage and intracellular Notch translocation to the nucleus to activate the transcription of downstream targets, including Hes1 and Hey1^16^. We performed quantitative RT-PCR and confirmed JM2 significantly decreased the expression of Notch downstream target Hes1 and Hey1 at the transcription level compared to antennapedia alone or JM2-scrambled in VTC-001 and VTC-037 GSCs (Figure 7F and G).

**Figure 7:**
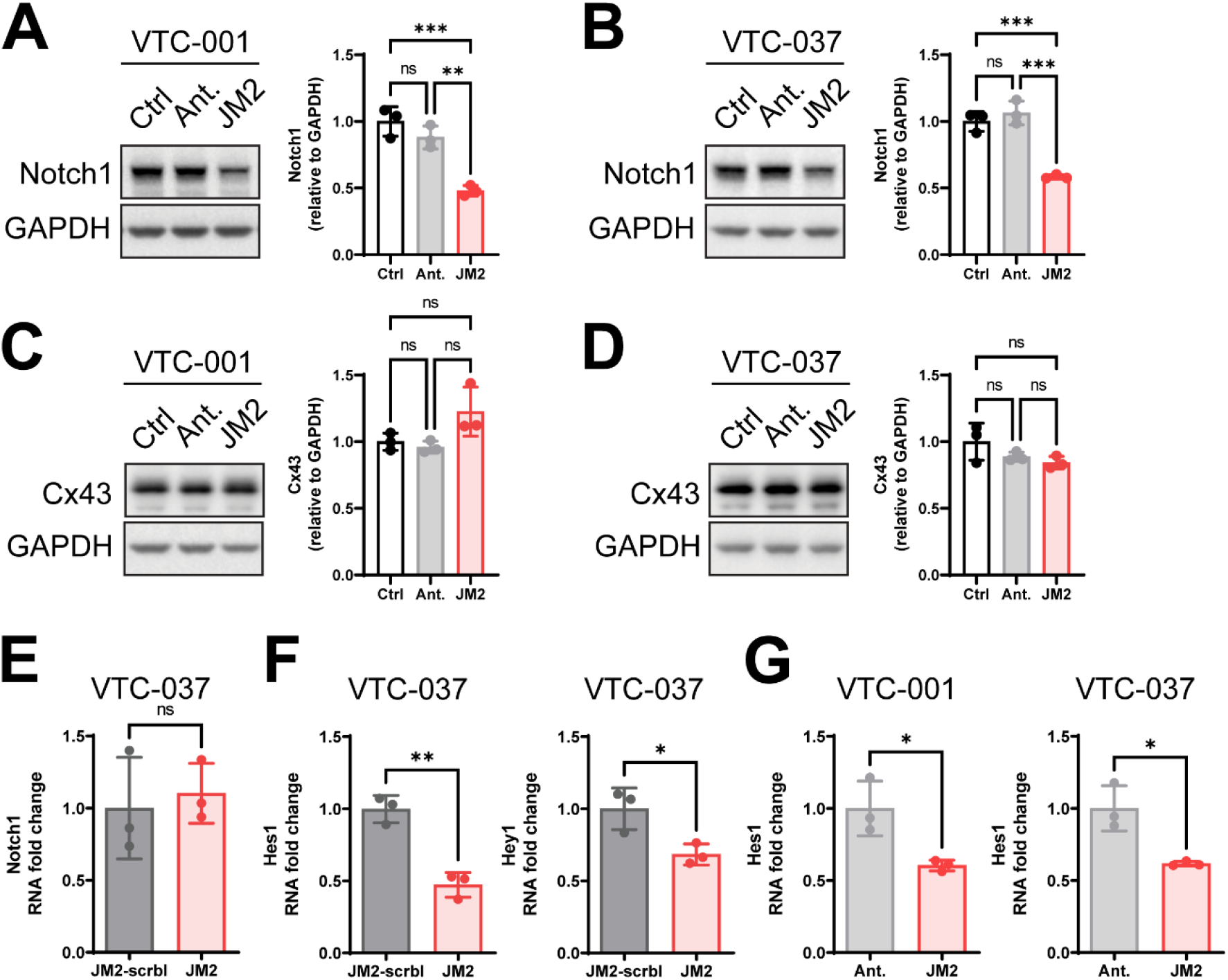
JM2 inhibits Notch signaling. VTC-001 GSCs and VTC-037 GSCs were treated or not with antennapedia (Ant.) or JM2 peptides at 50 μM for 24 h. Cell lysates were analyzed by immunoblotting using antibodies against Notch1 (**A-B** quantification shown on right), Cx43 (**C-D** quantification shown on right), and GAPDH as loading control. Statistical analysis was performed with one-way analysis of variance (ANOVA), with Tukey’s multiple comparisons test. \*\**p*≤0.01, \*\*\**p*≤0.001, ns: non significant. Data are represented as mean ± SEM. **E)** VTC-037 GSCs cells were treated or not with JM2-scrambled (JM2-scrbl) or JM2 peptides at 50 μM for 24 h, RNA was extracted and *Notch1* mRNA levels were quantified by qRT-PCR. **F)** VTC-037 GSCs cells were treated or not with JM2-scrambled (JM2-scrbl) or JM2 peptides at 50 μM for 24 h, RNA was extracted and *Hes1* and *Hey1* mRNA levels were quantified by qRT-PCR. **G)** VTC-001 and VTC-037 GSCs cells were treated or not with antennapedia (Ant.) or JM2 peptides at 50 μM for 24 h, RNA was extracted and *Hes1* mRNA levels were quantified by qRT-PCR. A two-tailed unpaired Student’s *t*-test was used; \**p*≤0.05, \*\**p*≤0.01. Data are represented as mean ± SEM.

### JM2 peptide decreases GSC-derived tumor growth in vivo

To test the effect of JM2 on GSCs *in vivo*, we utilized previously characterized GSCs isolated and enriched from human GBM LN229 cells (LN229 GSCs). These cells express Cx43 and form tumors following injection in mouse flank^21^. We first assessed the effect of JM2 on LN229 GSCs *in vitro*, and confirmed JM2 significantly decreased LN229 GSC survival after 4 days, similar to the results obtained with other GSC cell lines in this study (Figure 8A). Next, we injected LN229 GSCs in mouse flanks, and upon tumor formation 7 days later, we administered JM2 intratumorally every other day (Figure 8B). We observed a significant decrease in tumor growth in mice treated with JM2 compared to control or JM2-scrambled after 33 days (Figure 8C). Tumor sections were stained using TUNEL, and we observed increased LN229 GSC death in tumors treated with JM2 compared to controls (Figure 8D).

**Figure 8:**
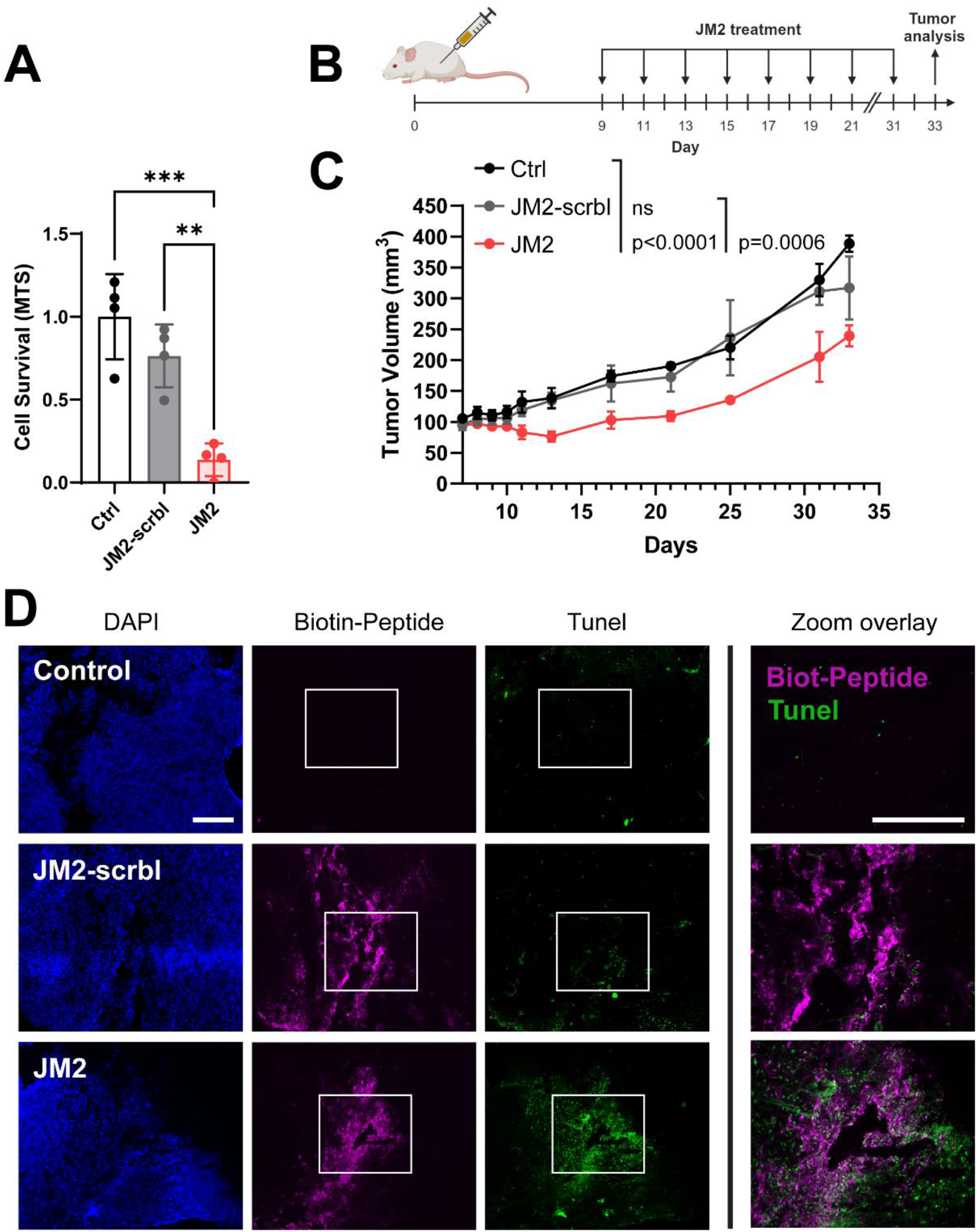
JM2 inhibits GSC survival *in vivo*. **A)** LN229 GSCs were treated or not with 100 μM of JM2-scrambled (JM2-scrbl) or JM2 peptides for 4 days before cell survival was assessed using an MTS assay. Statistical analysis was performed with one-way analysis of variance (ANOVA) with Tukey’s multiple comparisons test. \*\**p*≤0.01, \*\*\**p*≤0.001. Data are represented as mean ± SEM. **B)** LN229 GSCs were injected in mouse flank and upon tumor formation, JM2-scrambled (JM2-scrbl) or JM2 at 300 μM were administered intratumorally every other day (created with BioRender.com). **C)** Tumor volume was determined at different time points (n=3). The rate of tumor growth over the first 31 days was different among all three interventions (X2(2)=28.1845; p<0.0001). Specifically, the rate of tumor growth in JM2 was significantly slower than that of the Control (t(87)=-5.081; p<0.0001) and JM2-scrambled (t(87)=-3.873; p=0.0006). **D)** After 33 days, tumor sections from B were analyzed for the presence of biotin-tagged JM2-scrambled (JM2-scrbl) and JM2 using streptavidin-conjugated to AlexaFluor647 (magenta), and cell death was assessed using TUNEL staining (green). DAPI staining was used to detect nuclei (Scale bar: 400 μm).

## Discussion

The role of Cx43 in cancer progression is dynamic and complex, with Cx43 described as both a tumor suppressor and oncogenic protein, depending on cancer type and stage^18^. This is not only due to differential expression of Cx43 during cancer progression, but is also linked to channel-dependent and -independent functions of Cx43 and its roles in dynamic protein complex formation in regulation of cell proliferation, migration/invasion, and apoptosis^18,34,35^. Increased levels of Cx43 correlate with TMZ resistance in GBM cells, and GBM patients that present high levels of Cx43 mRNA and low tumor levels of O6-Methylguanine-DNA Methyltransferase (MGMT), an enzyme that repairs TMZ-induced DNA lesions, have significantly shorter life spans than those with low levels of Cx43 mRNA^19-21^. In addition, brain metastatic cancer cells utilize Cx43 gap junctions to communicate with normal astrocytes to support tumor growth, invasion, and chemoresistance via, among other mechanisms, the formation of a nanotube communication networks^22^. In differentiated GBM cell lines, expression of a dominant negative Cx43 mutant that blocked gap junction and cell-cell communication was found to increase cell invasion^36^. However, Cx43 is also associated with anti-proliferative effects in glioma and reduced levels of Cx43 protein was reported in high-grade gliomas^23,24^. Our study isolates a novel and specific cytosolic non-channel function of Cx43 in complexing with microtubules to promote GSC maintenance and survival, independent of other channel-based/tumor suppressive Cx43 functions.

GSCs play a critical role in GBM resistance to treatment and tumor recurrence^5-8^. Proliferating GSCs display a loss of Cx43-dependent gap junctions at the plasma membrane that is accompanied by reduced intercellular communication, and Cx43 has been shown to be expressed at much lower levels in GSCs compared to their differentiated counterparts^26,27^. Although we confirmed Cx43 expression is low in GSCs compared to differentiated GSCs, we demonstrate, for the first time, an enrichment of Cx43 non-junctional cytoplasmic localization at the microtubules in GSCs. While Cx43 is known to oligomerize at the trans-Golgi network before trafficking to the plasma membrane through vesicular transport along microtubules^25^, our implementation of stochastic resolution microcopy allowed us to parse out an intracellular population of Cx43 directly interacting with microtubules independent from the cytoplasmic Cx43 hemichannels being transported in vesicles along microtubules. Localizing these molecules at 25 nm resolution supersedes previous biochemical and imaging techniques, which failed to distinguish this novel phenomenon – namely, two functionally distinct cytoplasmic populations of Cx43 intimately associated with the microtubule cytoskeleton.

Microtubules are critical in controlling key cellular processes including division, motility, differentiation, transport of cargoes and vesicles, and overall survival of cells^37^. The functions, together with the stability, and highly dynamic properties of microtubules, are regulated by post-translational modifications and interactions with microtubule binding proteins (MTBPs)^38^. MTBPs present different roles in stabilizing/destabilizing microtubules, as well as in anchoring microtubules to cytoskeletal components. Here, we identify Cx43 as a novel MTBP in GSCs, however, the exact role of Cx43 in regulating microtubule function, as well as the precise subcellular compartment within which Cx43 interacts with microtubules, remain to be elucidated. In addition to modulating microtubule dynamics to affect vesicular transport and/or cell membrane and junctional remodeling, Cx43 complexing with tubulin may compete with signaling molecules to elicit activation/inactivation of pathways regulating survival and differentiation, for example.

The 14 amino acid tubulin-interacting domain within the ∼130 amino acid CT of Cx43 has been well characterized ^28,39^. One study identified residues ^239^RV^240^ and ^247^YHAT^250^ to be critical for Cx43 interaction with microtubules^40^. Furthermore, Src phosphorylation of Cx43 on Y^247^ has been shown to disrupt this interaction, suggesting a dynamic process of Cx43 interaction with microtubules^40^. The Cx43 interaction domain on tubulin, however, has not been well characterized. Using in vitro recombinant proteins, it has been reported that the domain that comprises residues 114-243 on β-tubulin is necessary for Cx43 to interact with microtubules^41^. Moreover, differential expression of tubulin in GBM CSCs has been speculated to impact resistance to chemotherapeutics, many of which historically target microtubules^42^. Our findings therefore indicate dynamic complexing of Cx43 with tubulin can be modulated by the cell and enhanced to maintain specific states advantageous to GBM progression, including in the maintenance of GSCs.

Using JM2, a Cx43 mimetic peptide of Cx43 tubulin-binding domain, we confirm efficient and specific disruption of Cx43-microtubule interaction in GSCs together with enhanced killing of this cancer stem cell population, highlighting a novel tumorigenic role for Cx43 in GSCs, and GBM. Thermal shift assays reveal JM2 presents high affinity for both α- and β- tubulin. While chemotherapeutic drugs such as paclitaxel have been used to target microtubule functions as therapeutic agents, many of these therapeutic agents cause significant side-effects^43^. Moreover, slower replicating cancer cells, including GSCs, are resistant to such strategies^44^. Here we highlight a novel strategy targeting Cx43/microtubule complexing alone, and presumably maintaining other roles of the cytoskeleton in normal tissues. It is possible that the Cx43 sub-population we identify as directly bound to microtubules is in the endoplasmic reticulum, potentially providing anchoring and stabilizing cytoskeletal ‘highways’ within the GSC that are disorganized by JM2. Prior work with JM2 indicated a loss of Cx43 delivery to the cell surface due to increased Cx43/tubulin complexing, but this work did not attempt to isolate vesicular Cx43 *vs* that directly bound to microtubules^31^. Additionally, the cellular context of primary GSCs (which have few gap junctions to begin with) further highlights the specificity of this biology to such cells.

Mimetic peptides of Cx43 have been implemented to efficiently modulate Cx43 channel-dependent and -independent functions^45^. The Cx43 mimetic peptide of the ZO-1 binding domain aCT1 that selectively inhibits Cx43 hemichannel activity and increases Cx43 dependent gap junction formation^46,47^, restores TMZ sensitivity to TMZ-resistant GBM cells^21^. A Cx43 mimetic peptide of a short region within Cx43 C-terminus that recruits and inhibits c-Src activity, decreases cell motility, invasion, and proliferation in GSCs^48^. Peptide-based cancer therapies have recently gained increasing recognition and validation in the field. In fact, peptides are presenting several advantages when compared to small molecules and kinase inhibitors with peptides demonstrating higher specificity in targeting cancer cells and/or their associated tumorigenic aspects, and exerting lower toxicity in normal tissues^49^. Importantly, although astrocytes express high level of Cx43, we observed no toxic effect of JM2 on normal human astrocytes *in vitro*, suggesting that Cx43/microtubule complex formation is cell-type specific in its putative role in GSC maintenance.

We further demonstrated that JM2 significantly decreased Notch expression and downstream signaling in GSCs, wherein we determined downregulation of *Hes1* and *Hey1* transcription in response to the Cx43 peptide. Activation of the Notch pathway is associated with GSC proliferation, self-renewal, and survival, and maintenance of stemness in GSCs^10,12^. Cell-cell contacts lead to interaction and activation of Notch receptors by Delta-like or Jagged ligands on apposing cells, prior to two enzymatic cleavages that promote internalization and nuclear translocation of the Notch intracellular domain (NICD). There, NICD regulates the transcription of genes such as *Hey1* and *Hes1*, belonging to the basic helix-loop-helix (bHLH) family of transcription factors, which in turn, regulate the expression of genes involved in the maintenance of stem-like properties in GSCs^12^. The vascular niche of GBM tumors has been shown to sustain GSC survival^50^. Given the major role of Notch signaling in the vasculature, high levels of Delta-like/Jagged ligands likely maintain GSCs in these niches. While we do not know the exact mechanism of JM2-mediated Notch disruption, alterations in microtubule trafficking likely dull GSC ability to engage with and respond to ligands, compromising survival.

Given the lack of progress in treatment of GBM, new approaches in targeting not just GBM cancer cells, but also GSC populations likely persisting after surgical resection of tumors, are critical. Based on our preclinical findings, administration of JM2 in combination with chemotherapy (TMZ) may improve patient survival though slowing of tumor recurrence. By targeting a specific function of Cx43, and not gap junctions (which are essential in many organ systems), Cx43 expression, or tubulin polymerization (e.g. Taxol), we anticipate fewer side effects based on our data in normal astrocytes, for example. While a promising class of drugs, peptides do present their challenges in the context of administration and low stability *in vivo*. Delivery systems such as nanoparticles show some potential here^51^. Extracellular vesicles (EVs) are also emerging as a promising and specific means of delivery payloads such as peptide drugs to specific cell-types, including tumors^52,53^. Our work identifies Cx43/tubulin complexing as a specific event in GSCs, and represents a step forward in clarifying oncogenic *vs* tumor suppressive roles for connexins in GBM. The therapeutic approach we posit, and provide data for, maintains the homeostatic functions of gap junctions, whilst selectively targeting Cx43/tubulin complexes with downstream perturbation of Notch signaling pathways required for GSC survival. In sum, JM2 presents a mechanistically intriguing and promising new GBM therapeutic for treating this devastating disease.

## Acknowledgments

The authors thank Jane Jourdan and Pratik Kanabur for technical support, and Dr. Allison N. Tegge (Fralin Biomedical Research Institute, Department of Statistics, Virginia Tech) for consultation on the statistical method used for tumor growth. This work was supported in part by a National Institutes of Health NCI R41 grant (CA217503 to S.L.) and NHLBI R35 (HL161237-02 to R.G.G.).

## Declaration of interests

S.L. is Co-founder and CEO of Acomhal Research Inc, which licensed the JM2 peptide. R.G.G. is Co-founder and CSO of Acomhal Research Inc. S.L. and R.G.G. have ownership interests in Acomhal Research Inc. The remaining authors have no disclosures to report.

---

**Supplemental Figure 1:**
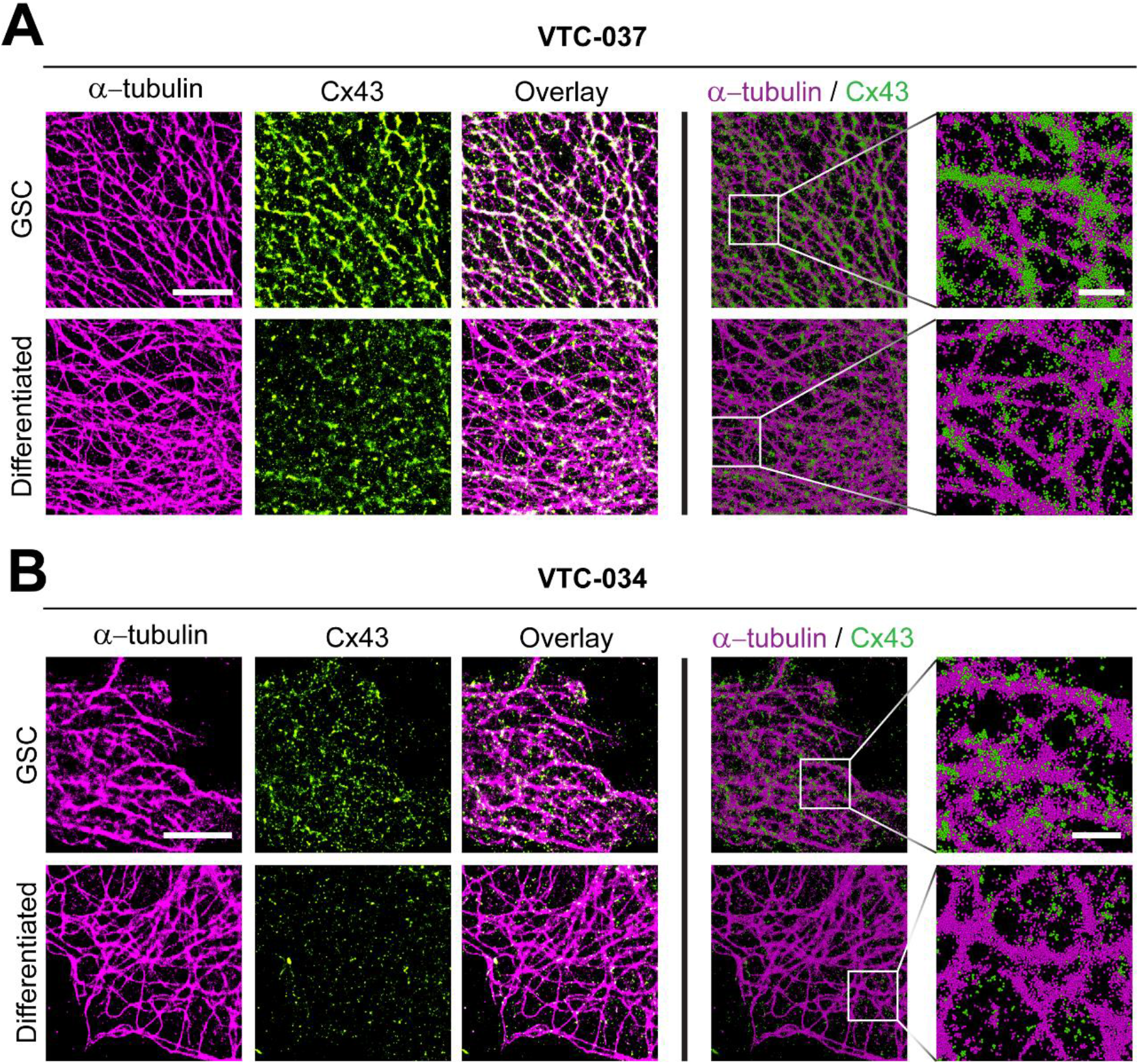
Increased Cx43 interaction with microtubules in GSCs. Stochastic optical reconstruction microscopy (STORM) derived localizations of Cx43 (green) and α-tubulin (magenta) in VTC-037 (**A**) or VTC-034 (**B**) GSCs or differentiated by addition of 10% FBS for 24 h. Left 6 panels - point-splatting is used to better identify co-localization (white; scale bar: 5 μm). Right 4 panels are point-clouds of 50 nm spheres representing individual localizations, including zoomed-in regions (scale bar: 1 μm).

**Supplemental Figure 2:**
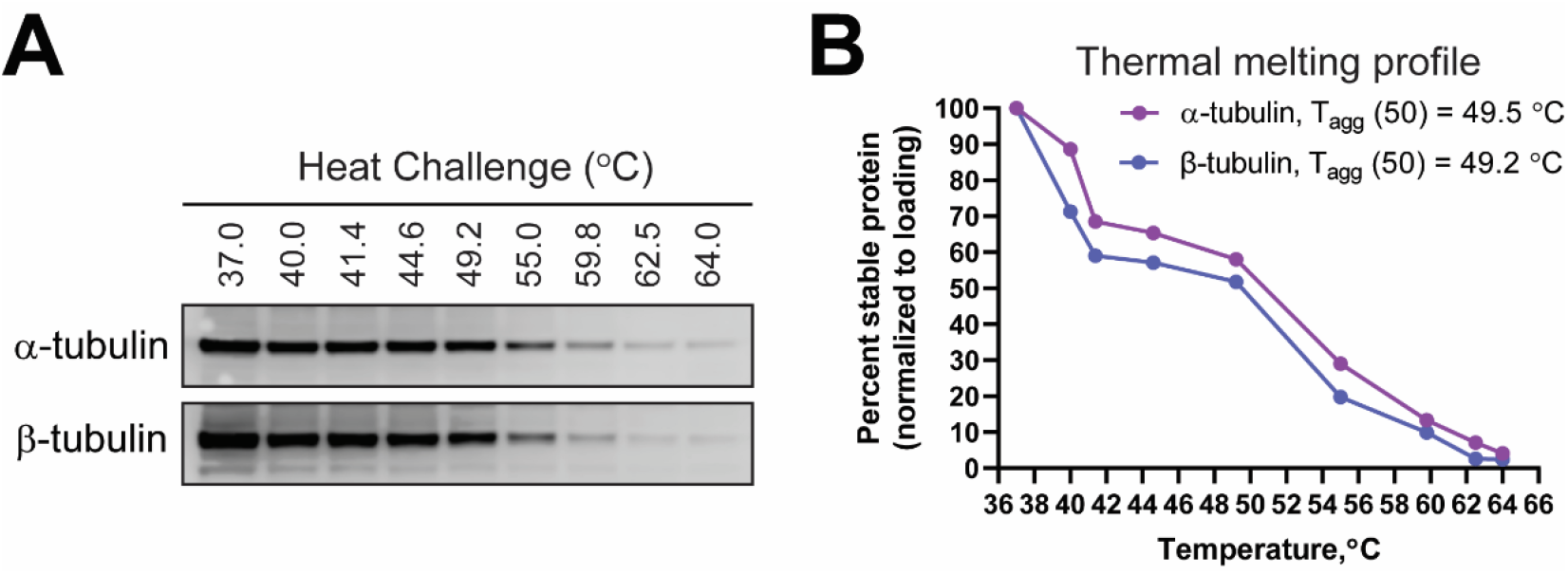
Cellular thermal shift assay in VTC-037 GSC lysates. Heat gradient was used on VTC-037 GSC lysates to determine thermal melting profiles of α- and β- tubulin, analyzed by immunoblotting (**A**), and represented as percent stable protein over increased temperatures (**B**).

